# Looking for a Signal in the Noise: Revisiting Obesity and the Microbiome

**DOI:** 10.1101/057331

**Authors:** Marc A Sze, Patrick D Schloss

## Abstract

Two recent studies have re-analyzed published data and found that when datasets are analyzed independently there was limited support for the widely accepted hypothesis that changes in the microbiome are associated with obesity. This hypothesis was reconsidered by increasing the number of datasets and pooling the results across the individual datasets. The Preferred Reporting Items for Systematic Reviews and Meta-Analyses (PRISMA) guidelines were applied to identify 10 studies for an updated and more synthetic analysis. Alpha diversity metrics and the relative risk of obesity based on those metrics were used to identify a limited number of significant associations with obesity; however, when the results of the studies were pooled using a random effects model significant associations were observed between Shannon diversity, number of observed OTUs, and Shannon evenness and obesity status. They were not observed for the ratio of *Bacteroidetes* and *Firmicutes*or their individual relative abundances. Although these tests yielded small P-values, the difference between the Shannon diversity index of non-obese and obese individuals was 2.07%. A power analysis demonstrated that only one of the studies had sufficient power to detect a 5% difference in diversity. When Random Forest machine learning models were trained on one dataset and then tested using the other 9 datasets, the median accuracy varied between 33.01 and 64.77% (median=56.68%). Although there was support for a relationship between the microbial communities found in human feces and obesity status, this association was relatively weak and its detection is confounded by large interpersonal variation and insufficient sample sizes.

**Importance:** As interest in the human microbiome grows there is an increasing number of studies that can be used to test numerous hypotheses across human populations. The hypothesis that variation in the gut microbiota can explain or be used to predict obesity status has received considerable attention and is frequently mentioned as an example for the role of the microbiome in human health. Here we assess this hypothesis using ten independent studies and find that although there is an association, it is smaller than can be detected by most microbiome studies. Furthermore, we directly tested the ability to predict obesity status based on the composition of an individual’s microbiome and find that the median classification accuracy is between 33.01 and 64.77%. This type of analysis can be used to design future studies and expanded to explore other hypotheses.

## Introduction

Obesity is a growing health concern with approximately 20% of the youth (aged 2–19) in the United States classified as either overweight or obese (1). This number increases to approximately 35% in adults (aged 20 or older) and these statistics have seen little change since 2003 (1). Traditionally, the body mass index (BMI) has been used to classify individuals as non-obese or obese (2). Recently, there has been increased interest in the role of the microbiome in modulating obesity (3, 4). If the microbiome does affect obesity status, then manipulating the microbiome could have a significant role in the future treatment of obesity and in helping to stem the current epidemic.

There have been several studies that report observing a link between the composition of microbiome and obesity in animal models and in humans. The first such study used genetically obese mice and observed the ratio of the relative abundances of *Bacteroidetes* to *Firmicutes* (B:F) was lower in obese mice than lean mice (5). Translation of this result to humans by the same researchers did not observe this effect, but did find that obese individuals had a lower alpha-diversity than lean individuals (6). They also showed that the relative abundance of *Bacteroidetes* and *Firmicutes* increased and decreased, respectively, as obese individuals lost weight while on a fat or carbohydrate restricted diet (7). Two re-analysis studies by Walters et al. (8) and Finucane et al. (9) interrogated previously published microbiome and obesity data and concluded that the previously reported differences in community diversity and B:F among non-obese and obese individuals could not be generalized. Regardless of the results using human populations, studies using animal models where the community was manipulated with antibiotics or established by colonizing germ-free animals with varied communities appear to support the association since these manipulations yielded differences in animal weight (10–13). The purported association between the differences in the microbiome and obesity have been widely repeated with little attention given to the lack of a clear signal in human cohort studies.

The recent publication of additional studies that collected BMI data for each subject as well as other studies that were not included in the earlier re-analysis studies offered the opportunity to revisit the question relating the structure of the human microbiome to obesity. One critique of the prior re-analysis studies is that the authors did not aggregate the results across studies to increase the effective sample size. It is possible that there were small associations within each study that were not statistically significant because the individual studies lacked sufficient power. Alternatively, diversity metrics may mask the appropriate signal and it is necessary to measure the association at the level of microbial populations. The Walters re-analysis study demonstrated that Random Forest machine learning models were capable of predicting obesity status within a single cohort, but did not attempt to test the models on other cohorts. The purpose of this study was to perform a meta-analysis of the association between differences in the microbiome and obesity status by analyzing and applying a more systematic and synthetic approach than was used previously.

## Results

### Literature Review and Study Inclusion

To perform a robust meta-analysis and limit inclusion bias, we followed the Preferred Reporting Items for Systematic Reviews and Meta-Analyses (PRISMA) guidelines to identify the studies that we analyzed (14). A detailed description of our selection process and the exact search terms are provided in the Supplemental Text and in Figure 1. Briefly, we searched PubMed for original research studies that involved studying obesity and the human microbiome. The initial search yielded 187 studies. We identified 10 additional studies that were not designed to explicitly test for an association between the microbiome and obesity. We then manually curated the 197 studies to select those studies that included BMI and 16S rRNA gene sequence data. This yielded 11 eligible studies. An additional study was removed from our analysis because no individuals in the study had a BMI over 30. Among the final 10 studies, 3 were identified from our PubMed search (10, 15, 16), 5 were originally identified from the 10 studies that did not explicitly investigate obesity but included BMI data (17–21), and two datasets were used (22, 23) because these publications did not specifically look for any metabolic or obesity conditions but had control populations and enabled us to help mitigate against publication biases associated with the bacterial microbiome and obesity. The ten studies are summarized in Table 1. For comparison, two of these studies were included in the Finucane re-analysis study (10, 21) and four of these studies were included in the Walters re-analysis study (10, 15, 20, 21). The 16S rRNA gene sequence data from each study was re-analyzed using a similar approach based on previously described methods for reducing the number of chimeric sequences and sequencing errors for 454 and Illumina MiSeq data (24, 25). The sequences were clustered into operational taxonomic units (OTUs) using the average neighbor approach (26) and into taxonomic groupings based on their classification using a naive Bayesian classifier (27).

**Figure 1:**
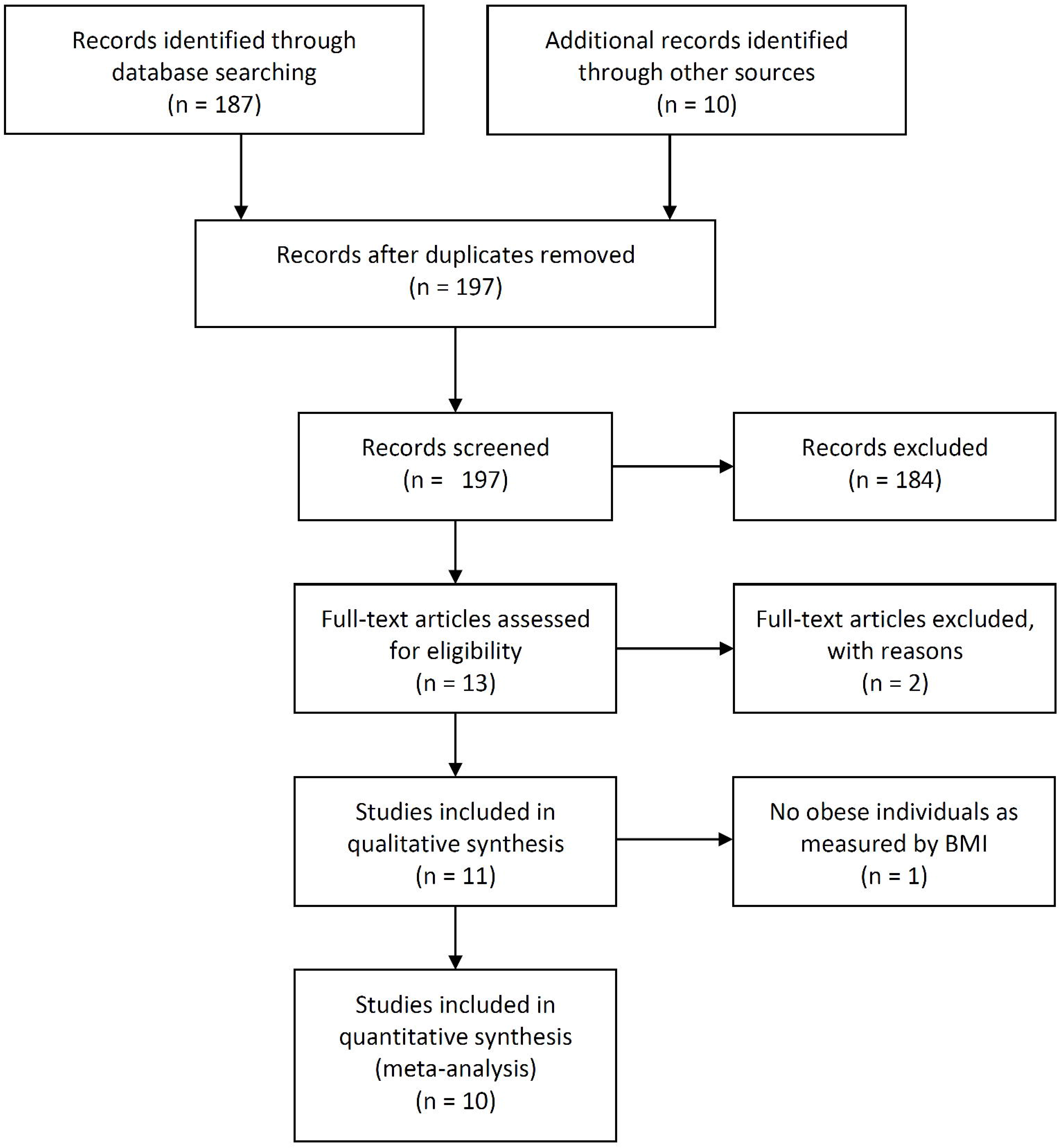
PRISMA flow diagram of total records searched (39).

**Table 1.**
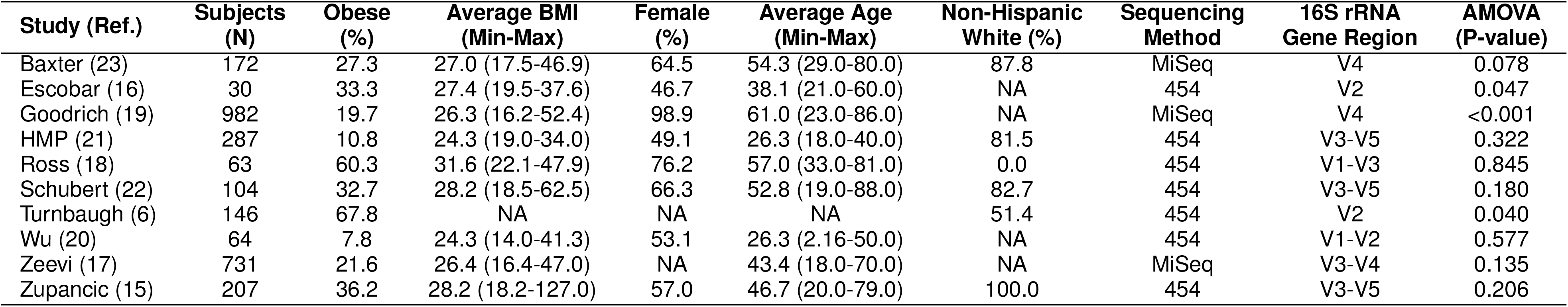
Summary of obesity, demographic, sequencing, and beta-diversity analysis data for the studies used in the meta-analysis. NA indicates that those metadata were not available for that study.

### Alpha diversity analysis

We calculated the Shannon diversity index, observed richness, and Shannon evenness, the relative abundance of *Bacteroidetes* and *Firmicutes,* and the ratio of their relative abundance (B:F) for each sample. Once we transformed each of the six alpha diversity metrics to make them normally distributed, we used a t-test to identify significant associations between the alpha diversity metric and whether an individual was obese for each of the ten studies. The B:F and the relative abundance of *Firmicutes* were not significantly associated with obesity in any study. We identified seven P-values that were less than 0.05: three studies indicated obese individuals had a lower richness, two studies indicated a significantly lower diversity, one study indicated a significantly lower evenness, and one study indicated a significantly higher relative abundance of *Bacteroidetes* (Figures 2 and S1). These results largely match those of the Walters and Finucane re-analysis studies. Interestingly, although only two of the ten studies observed the previously reported association between lower diversity and obesity, the other studies appeared to have the same trend, albeit the differences were not statistically significant. We used a random effects linear model to combine the studies using the study as the random effect and found statistical support for decreased richness, evenness, and diversity among obese individuals (all P<0.011). Although there was a significant relationship between these metrics and obesity status, the effect size was quite small. The obese individuals averaged 7.47% lower richness, 0.88% lower evenness, and 2.07% lower diversity. There were no significant associations when we pooled the phylum-level metrics across studies. These results indicate that obese individuals do have a statistically significant lower diversity than non-obese individuals; however, it is questionable whether the difference is biologically significant.

**Figure 2:**
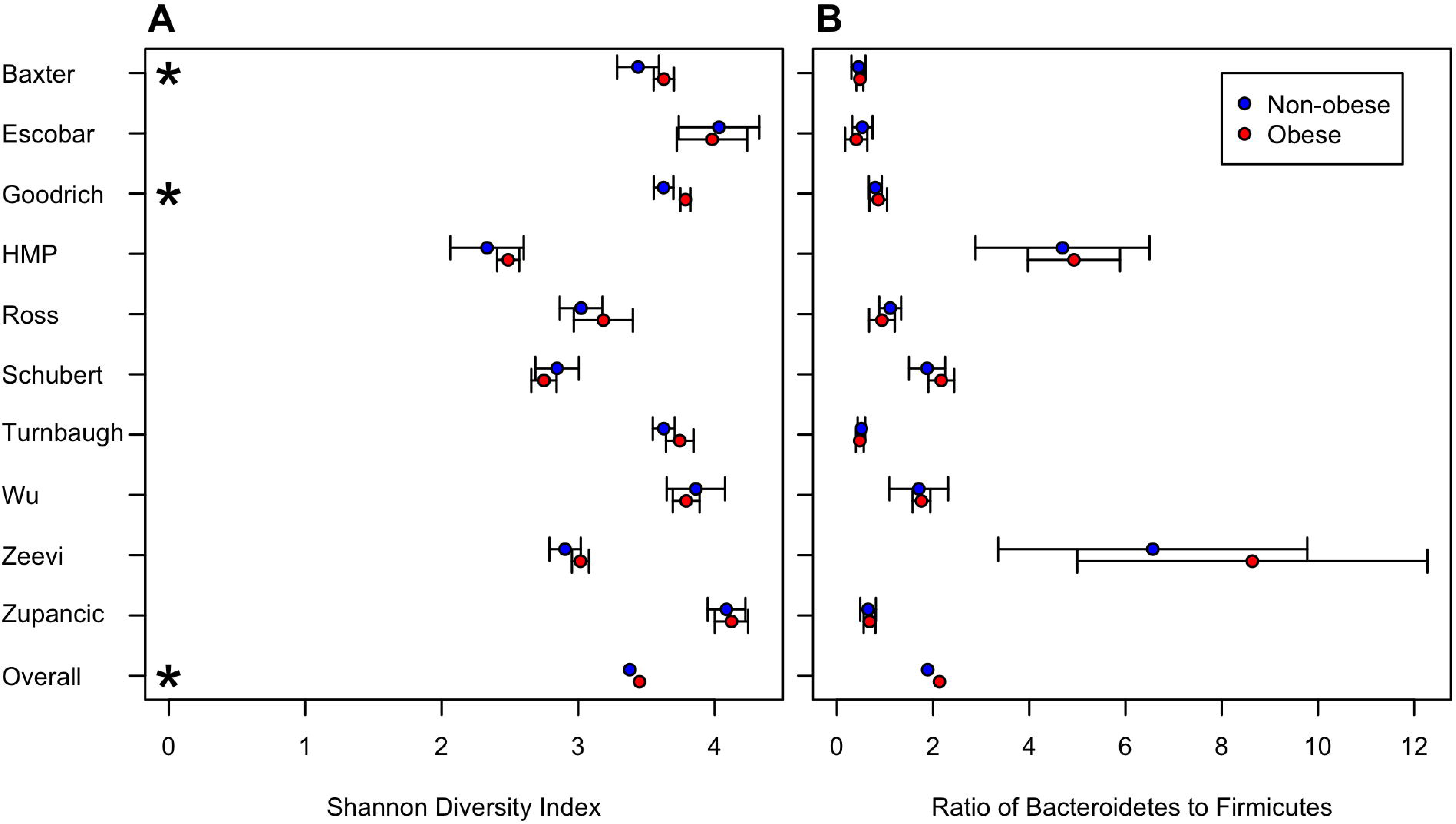
Individual and combined comparison of obese and non-obese groups for Shannon diversity (A) and B:F (B).

### Relative risk

Building upon the alpha diversity analysis we calculated the relative risk of being obese based on an individual’s alpha diversity metrics relative to the median metric for that study. Inspection of funnel plots for each of the metrics suggested that the studies included in our analysis were not biased (Figure S2). The results using relative risk largely matched those of using the raw alpha diversity data. Across the ten studies and six metrics, the only significant relative risk values were the richness, evenness, and diversity values from the Goodrich study (Figures 3 and S3). Again, although the relative risk values were not significant for other studies, the values tended to be above one. When we pooled the data using a random effects model, the relative risk associated with having a richness, evenness, or diversity below the median for the population was significantly associated with obesity (all P<0.0044). The relative risks associated with alpha diversity were small. The relative risk of having a low richness was 1.30 (95% CI: 1.13-1.49), low evenness was 1.20 (95% CI: 1.06-1.37), and low diversity was 1.27 (95% CI: 1.09-1.48). There were no significant differences in the phylum-level metrics. Again, the relative risk results indicate that individuals with a lower richness, evenness, or diversity are at statistically significant increased risk of being obese, it is questionable whether that risk is biologically or clinically relevant.

**Figure 3:**
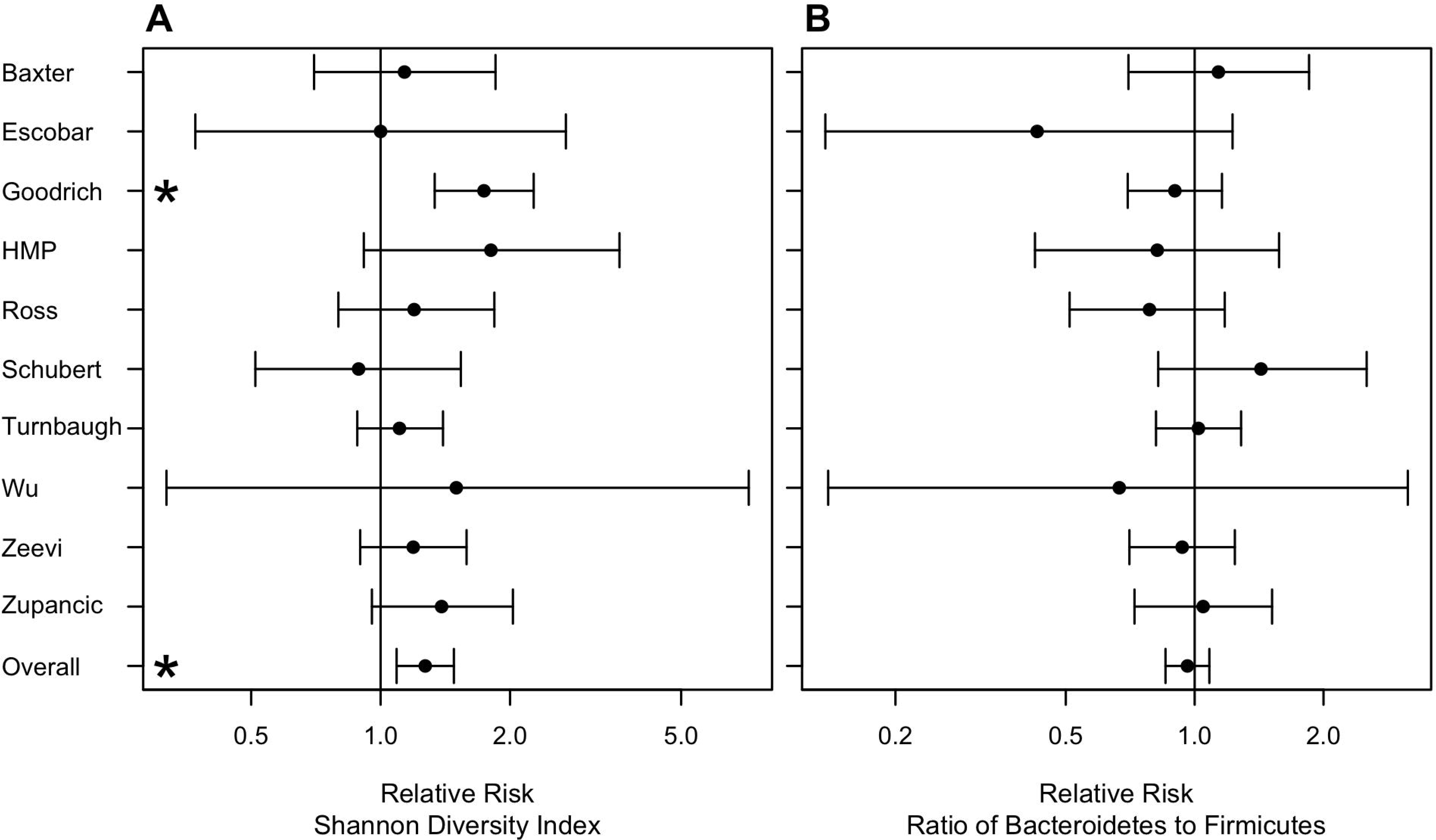
Meta analysis of the relative risk of obesity based on Shannon diversity (A) or B:F(B).

### Beta diversity analysis

Following the approach used by the Walters and Finucane re-analysis studies, for each dataset we calculated a Bray-Curtis distance matrix to measure the difference in the membership and structure of the individuals from each study. We then used AMOVA to test for significant differences between the structure of non-obese and obese individuals (Table 1). The Escobar, Goodrich, and Turnbaugh datasets indicated a significant difference in community structure (all P<0.05). Because it was not possible to ascertain the directionality of the difference in community structure because the samples are arrayed in a non-dimensional space or perform a pooled analysis using studies that had non-overlapping 16S rRNA gene sequence regions, it is unclear whether these differences reflect a broader, but perhaps small, shift in community structure between non-obese and obese individuals.

### Development of a microbiome-based classifier of obesity

The Walters re-analysis study suggested that it was possible to classify individuals as being non-obese or obese based on the composition of their microbiota. We repeated this analysis with additional datasets using OTU and genus-level phylotype data. For each study we developed a Random Forest machine learning model to classify individuals. Using ten-fold cross validation, the cross-validated AUC values for the OTU-based models varied between 0.52 and 0.69 indicating a relatively poor ability to classify individuals (Figure 4A). To test models on other datasets, we trained models using genus-level phylotype data for each dataset. The cross-validated AUC values for the models applied to the training datasets varied between 0.51 and 0.65, again indicating a relatively poor ability to classify individuals from the original dataset (Figure 4B). For each model we identified the probability where the sum of the sensitivity and specificity was the highest. We then used this probability to define a threshold for calculating the accuracy of the models when applied to the other nine datasets (Figure 5). Although there was considerable variation in accuracy values for each model, the median accuracy for each model varied between 0.33 (Turnbaugh) and 0.65 (HMP) (median=0.57). We built similar models using taxonomic representation based on phylum, class, order, and family assignments and saw no improvement in the results (Figure S4). We also attempted to predict individual BMI values as continuous variables based on the relative abundance of OTUs and genera. The median percent of the variance explained with the resulting models was 12.9% for the OTU-based models and 8.2% for the genus-based models. When we considered the number of samples, balance of non-obese and obese individuals, and region within the 16S rRNA gene for each study it was not possible to identify factors that predictably affected model performance. The ability to predict obesity status using relative abundance data from the communities was only marginally better than random. These results suggest that given the large diversity of microbiome compositions it is difficult to identify a taxonomic signal that can be associated with obesity.

**Figure 4:**
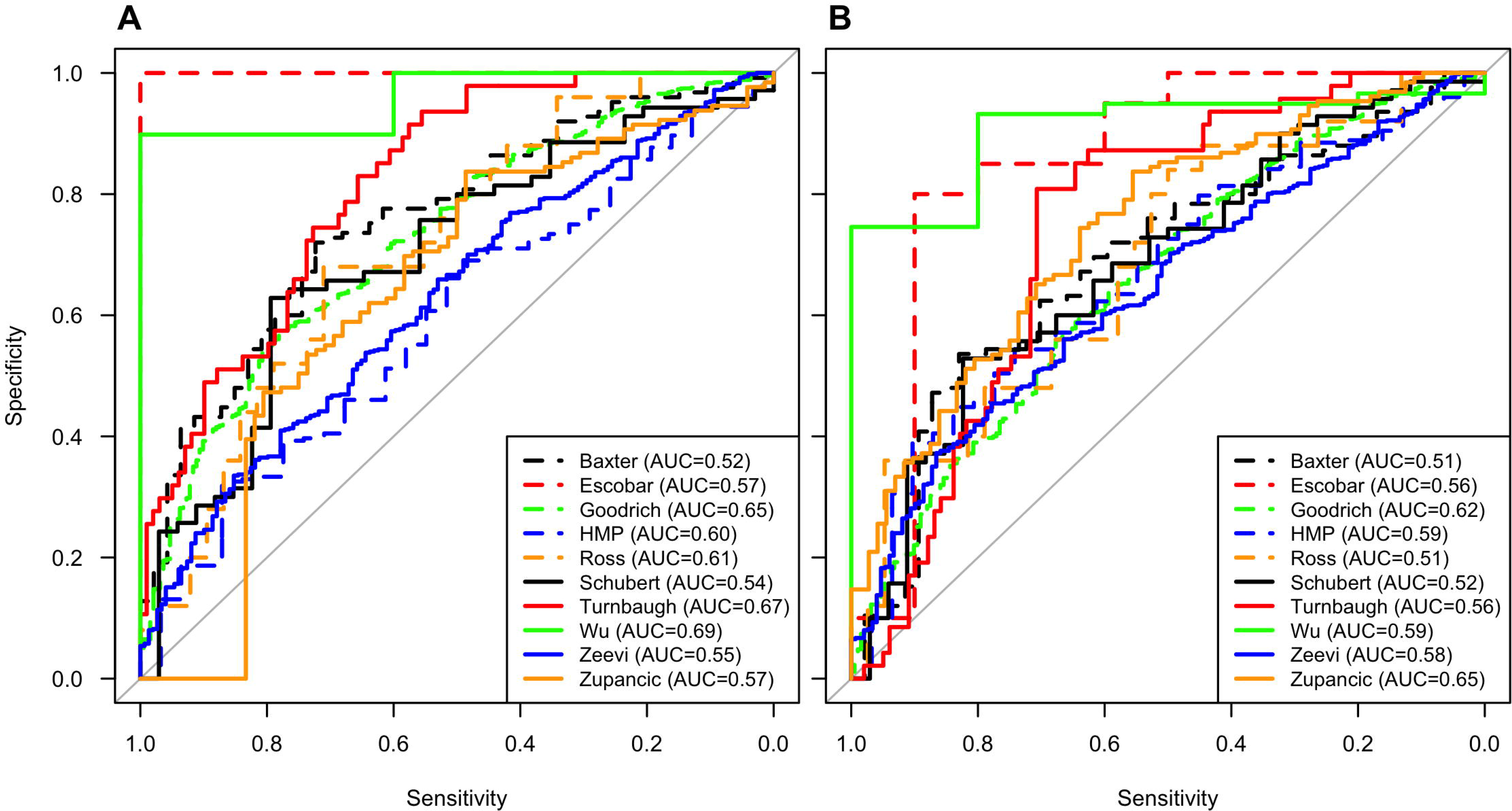
ROC curves for each study based on classification of non-obese or obese groups using OTUs (A) or genus-level classification (B).

**Figure 5:**
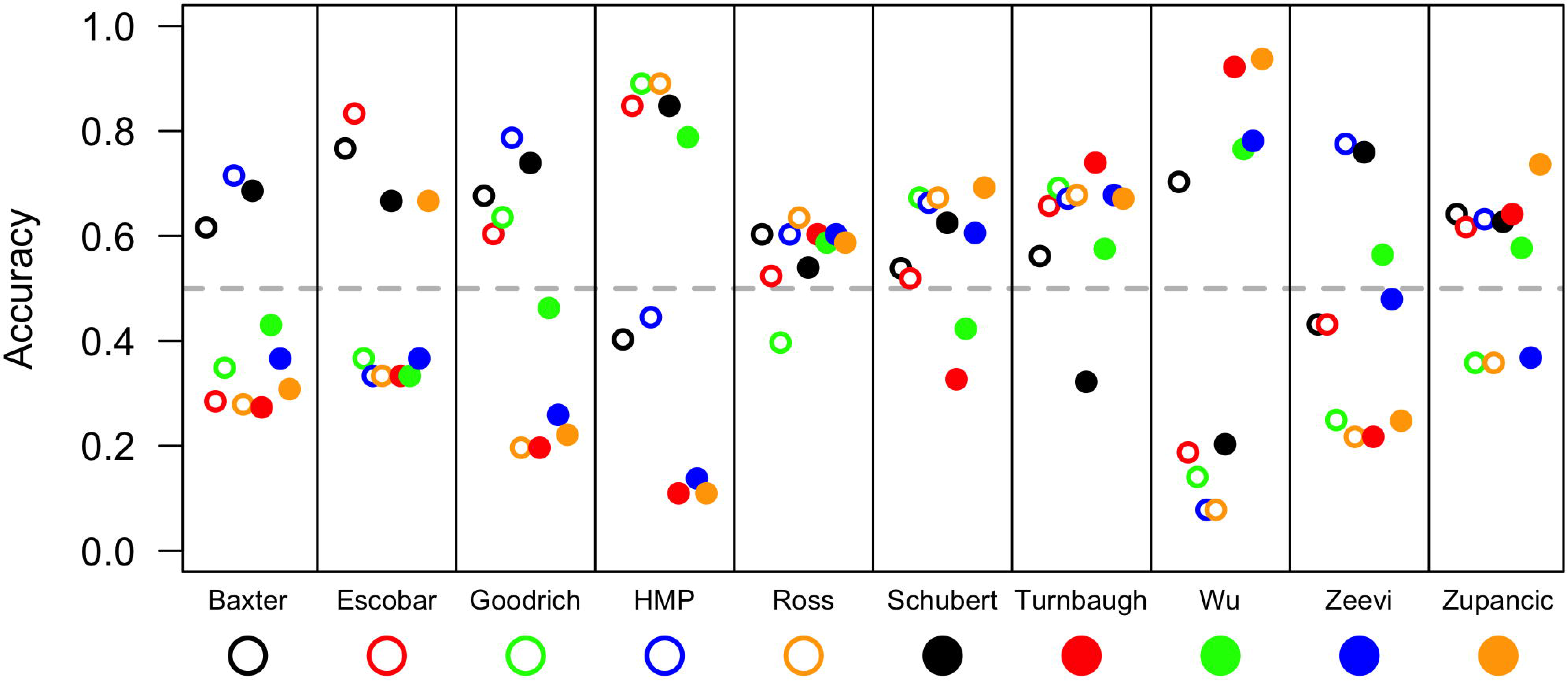
Overall accuracy of each study to predict non-obese and obese individuals based on that study’s Random Forest machine learning model applied to each of the other studies.

### Power and Sample Size Estimate Simulations

The inability to detect a difference between non-obese and obese individuals could be due to the lack of a true effect or because the study had insufficient statistical power to detect a difference because of insufficient sampling, large interpersonal variation, or unbalanced sampling of non-obese and obese individuals. To assess these factors, we calculated the power to detect differences of 1, 5, 10, and 15% in each of the alpha diversity metrics using the sample sizes used in each of the studies (Figures 6, S5–S10). Although there is no biological rationale for these effect sizes, they represent a range that includes effect sizes that would be generally considered to be biologically significant. Only the Goodrich study had power greater than 0.80 to detect a 5% difference in Shannon diversity and six of the studies had enough power to detect a 10% difference (Figure 6A). None of the studies had sufficient power to detect a 15% difference between B:F values (Figure S5). In fact, the maximum power among any of the studies to detect a 15% difference in B:F values was 0.25. Among the tests for relative risk, none of the studies had sufficient power to detect a Cohen’s d of 0.10 and only two studies had sufficient power to detect a Cohen’s d of 0.15. We next estimated how many individuals would need to have been sampled to have sufficient power to detect the four effect sizes assuming the observed interpersonal variation from each study and balanced sampling between the two groups (Figure 6B). To detect a 1, 5, 10, or 15% difference in Shannon diversity, the median required sampling effort per group was approximately 3,400, 140, 35, or 16 individuals, respectively. To detect a 1, 5, 10, and 15% difference in B:F values, the median required sampling effort per group was approximately 160,000, 6,300, 1,600, or 700 individuals, respectively. To detect a 1, 5, 10, and 15% difference in relative risk values using Shannon diversity, the median required sampling effort per group was approximately 39,000, 1,500, 380, or 170 individuals, respectively. These estimates indicate that most microbiome studies are underpowered to detect modest effect sizes using either metric. In the case of obesity, the studies were underpowered to detect the 0.90 to 6% difference in diversity that was observed across the studies.

**Figure 6:**
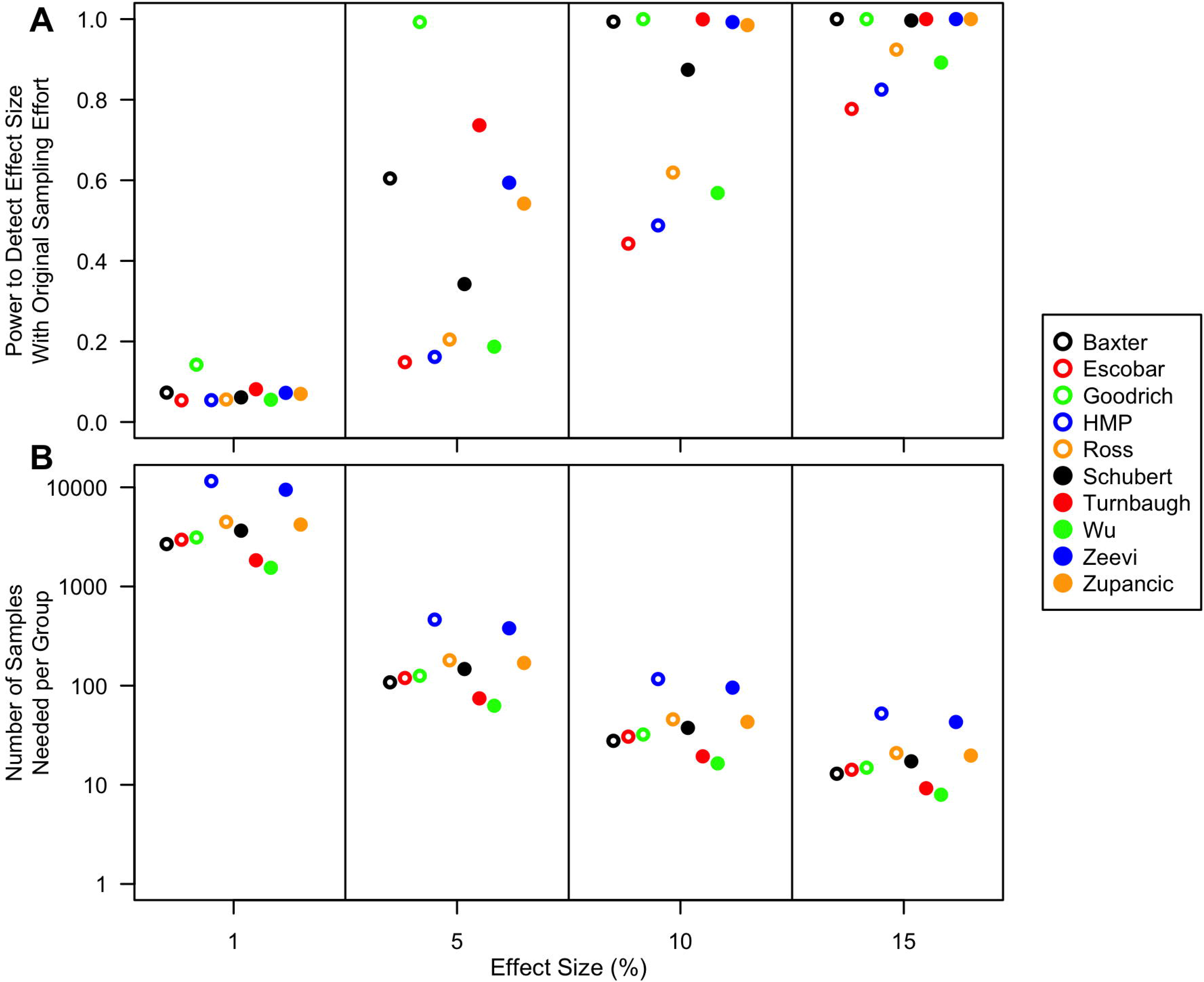
Power (A) and sample size simulations (B) for Shannon diversity for differentiating between non-obese versus obese for effect sizes of 1, 5, 10, and 15%. Power calculations use the sampling distribution from the original studies and the sample size estimations assume an equal amount of sampling from each treatment group.

## Discussion

Our meta-analysis helps to provide clarity to the ongoing debate of whether or not there are specific microbiome-based markers that can be associated with obesity. We performed an extensive literature review of the existing studies on the microbiome and obesity and performed a meta-analysis on the studies that remained based on our inclusion and exclusion criteria. By statistically pooling the data from ten studies, we observed significant, but small, relationships between richness, evenness, and diversity and obesity status as well as the relative risk of being obese based on these metrics. We also generated Random Forest machine learning models trained on each dataset and tested on the remaining datasets. This analysis demonstrated that the ability to reliably classify individuals as being obese based solely on the composition of their microbiome was limited. Finally, we assessed the ability of each study to detect defined differences in alpha diversity and observed that most studies were underpowered to detect modest effect sizes. Considering these datasets are among the largest published, it appears that most human microbiome studies are underpowered to detect differences in alpha diversity.

Alpha diversity metrics are attractive because they distill a complex dataset to a single value. For example, Shannon diversity is a measure of the entropy in a community and integrates richness and evenness information. Two communities with little taxonomic similarity can have the same diversity. Among ecologists the relevance of these metrics is questioned because it is difficult to ascribe a mechanistic interpretation to their relationship with stability or disease. Regardless, the concept of a biologically significant effect size needs to be developed among microbiome researchers. Alternative metrics could include the ability to detect a defined difference in the relative abundance of an OTU representing a defined relative abundance. What makes for a biologically significant difference or relative abundance is an important point that has yet to be discussed in the microbiome field. The use of operationally defined effect sizes should be adequate until it is possible to decide upon an accepted practice.

By selecting a range of possible effect sizes, we were able to demonstrate that most studies are underpowered to detect modest differences in alpha diversity metrics and phylum-level relative abundances. Several factors interact to limit the power of microbiome studies. There is wide interpersonal variation in the diversity and structure of the human microbiome. Some factors such as relationship between subjects could potentially decrease the amount variation (6) and other factors such as whether one lives in a rural environment could increase the amount of variation (28). In addition, the common experimental designs limit their power. As we observed, most of the studies included in our analysis were unbalanced for the variable that we were interested in. This was also true of those studies that originally sought to identify associations with obesity. Even with a balanced design, we showed that it was necessary to obtain approximately 140 and 6,300 samples per group to detect a 5% difference in Shannon diversity or B:F, respectively. It was interesting that these sample sizes agreed across studies regardless of their sequencing method, region within the 16S rRNA gene, or subject population (Figure 6). This suggests that regardless of the treatment or category, these sample sizes represent a good starting point for subject recruitment when using stool samples. Unfortunately, few studies have been published with this level of subject recruitment. This is troubling since the positive predictive rate of a significant finding in an underpowered study is small leading to results that cannot be reproduced (29). Future microbiome studies should articulate the basis for their experimental design.

Two previous re-analysis studies have stated that there was not a consistent association between alpha diversity and obesity (8, 9); however, neither of these studies made an attempt to pool the existing data together to try and harness the additional power that this would give and they did not assess whether the studies were sufficiently powered to detect a difference. Additionally, our analysis used 16S rRNA gene sequence data from ten studies whereas the Finucane study used 16S rRNA gene sequence data from three studies (7, 10, 21) and a metagenomic study (30) and the Walters study used 16S rRNA gene sequence data from five studies (10, 15, 20, 21, 28); two studies were included in both analyses (10, 21). Our analysis included four of these studies (10, 15, 20, 21) and excluded three of the studies because they were too small (7), only utilized metagenomic data (30), or used short single read Illumina HiSeq data that has a high error rate making it intractable for *de novo* OTU clustering (28). The additional seven datasets were published after the two reviews were performed and include datasets with more samples than were found in the original studies. Our collection of ten studies allowed us to largely use the same sequence analysis pipeline for all datasets and relied heavily on the availability of public data and access to metadata that included variables beyond the needs of the original study. To execute this analysis, we created an automated data analysis pipeline, which can be easily updated to add additional studies as they become available (https://github.com/SchlossLab/Sze_Obesity_mBio_2016/). Similarly, it would be possible to adapt this pipeline to other body sites and treatment or variables (e.g. subject’s sex or age).

Similar to our study, the Walters study generated Random Forest machine learning models to differentiate between non-obese and obese individuals (8). They obtained similar AUC values to our analysis; however, they did not attempt to test these models on the other studies in their analysis. When we performed the inter-dataset cross validation the median accuracy across datasets was only 56.68% indicating that the models did a poor job when applied to other datasets. This could be due to differences in subject populations and methods. Furthermore, others have reported improved classification at broader taxonomic levels (31); we did not find this to be the case across the studies in our analysis (Figure S4). Considering the median AUC for models trained and tested on the same data with ten-fold cross validation only varied between 0.51 and 0.65 and that there was not a strong signal in the alpha diversity data, we suspect that there is insufficient signal to reliably classify individuals to a BMI category based on their microbiota.

Although we failed to find an effect this does not necessarily mean that there is no role for the microbiome in obesity. There is strong evidence in murine models of obesity that the microbiome and level of adiposity can be manipulated via genetic manipulation of the animal and manipulation of the community through antibiotics or colonizing germ free mice with diverse fecal material from human donors (5, 10–13). These studies appear to conflict with the observations using human subjects. Recalling the large interpersonal variation in the structure of the microbiome, it is possible that each individual has their own signatures of obesity. Alternatively, it could be that the involvement of the microbiome in obesity is not apparent based on the taxonomic information provided by 16S rRNA gene sequence data. Rather, the differences could become more apparent at the level of a common set of gene transcripts or metabolites that can be produced from different structures of the microbiome.

## Methods

### Sequence Analysis Pipeline

All sequence data were publicly available and were downloaded from the NCBI Sequence Read Archive, the European Nucleotide Archive, or the investigators’ personal website (https://gordonlab.wustl.edu/TurnbaughSE/_10/_09/STM/_2009.html). In total seven studies used 454 (6, 15, 16, 18, 20–22) and three studies used Illumina sequencing (17, 19, 23). All of these studies used amplification-based 16S rRNA gene sequencing. Among the studies that sequenced the 16S rRNA gene, the researchers targeted the V1-V2 (20), V1-V3 (15, 16, 18), V3-V5 (21, 22), V4 [(19); (23);], and V3-4 (17) regions. For those studies where multiple regions were sequenced, we selected the region that corresponded to the largest number of subjects (6, 21). We processed the 16S rRNA gene sequence data using a standardized mothur pipeline. Briefly, our pipelines attempted to follow previously recommended approaches for 454 and Illumina sequencing data (24, 25). All sequences were screened for chimeras using UCHIME and assigned to operational taxonomic units (OTUs) using the average neighbor algorithm using a 3% distance threshold (26, 32). All sequence processing was performed using mothur (v.1.37.0) (33).

### Data Analysis

We split the overall meta-analysis into three general strategies using R (3.3.0). First, we followed the approach employed by Finucane et al (9) and Walters et al (8) where each study was re-analyzed separately to identify associations between BMI and the relative abundance of *Bacteroidetes* and Firmicutes, the ratio of *Bacteroidetes* and *Firmicutes* relative abundances (B:F), Shannon diversity, observed richness, and Shannon evenness (34). After each variable was transformed to fit a normal distribution a two-tailed t-test was performed for comparison of non-obese and obese individuals (i.e. BMI > 35.0). We performed a pooled analysis on these measured variables using linear random effect models to correct for study effect to asses differences on the combined dataset between non-obese and obese groups using the lme4 (v.1.1-12) R package (35). Next, we compared the community structure from non-obese and obese individuals using analysis of molecular variance (AMOVA) with Bray-Curtis distance matrices (36). This analysis was performed using the vegan (v.2.3–5) R package. For both analyses, the datasets were rarefied (N=1000) so that each study had the same number of sequences. Second, for each study we partitioned the subjects into a low or high group depending on whether their alpha diversity metrics were below or above the median value for the study. The relative risk (RR) was then calculated as the ratio of the number of obese individuals in the low group to the number of obese individuals in the high group. We then performed a Fisher exact-test to investigate whether the RR was significantly different from 1.0 within each study and across all of the studies using the epiR (0.9–77) and metafor (1.9–8) packages. Third, we used the AUCRF (1.1) R package to generate Random Forest models (37). For each study we developed models using either OTUs or genus-level phylotypes. The quality of each model was assessed by measuring the area under the curve (AUC) of the Receiver Operating Characteristic (ROC) using ten-fold cross validation. Because the genus-level phylotype models were developed using a common reference, it was possible to use one study’s model (i.e. the training set) to classify the samples from the other studies (i.e. the testing sets). The optimum threshold for the training set was set as the probability threshold that had the highest combined sensitivity and specificity. This threshold was then used to calculate the accuracy of the model applied to the test studies. To generate ROC curves and calculate the accuracy of the models we used the pROC (1.8) R package (38). Finally, we performed power and sample number simulations for different effect sizes for each study using the pwr (1.1–3) R package and base R functions. We also calculated the actual sample size needed based on the effect size of each individual study.

### Reproducible methods

A detailed and reproducible description of how the data were processed and analyzed can be found at https://github.com/SchlossLab/Sze_Obesity_mBio_2016/.

## Acknowledgements

The authors would like to thank Nielson Baxter and Shawn Whitefield for their suggestions on the development of the manuscript. We are grateful to the authors of the studies used in our meta-analysis who have made their data publicly available or available to us directly. Without their forethought studies such as this would not be possible. This work was supported in part by funding from the National Institutes of Health to PDS (U01AI2425501 and P30DK034933).

---

**Figure S1: Individual and Combined comparison of Obese and Non-Obese groups Based on Evenness (A), Richness (B), or the Relative Abundance of *Bacteroidetes* (C) and Firmictues (D)**.

**Figure S2: Funnel plots depicting the general lack of bias in the selection of datasets included in the analysis**.

**Figure S3: Meta Analysis of the Relative Risk of Obesity Based on Evenness (A), Richness (B), or the Relative Abundance of *Bacteroidetes* (C) and Firmictues (D)**.

**Figure S4: Overall accuracy of each study to predict non-obese and obese individuals based on that study’s Random Forest machine learning model applied to each of the other studies when trained using relative abundance of each phylum, class, order, family, or genus**. The cross-validated AUC values for the training model are provided for each study and taxonomic level.

**Figure S5: Power (A) and sample size simulations (B) for B:F for differentiating between non-obese versus obese for effect sizes of 1, 5, 10, and 15%**. Power calculations use the sampling distribution from the original studies and the sample size estimations assume an equal amount of sampling from each treatment group.

**Figure S6: Power (A) and sample size simulations (B) for richness for differentiating between non-obese versus obese for effect sizes of 1, 5, 10, and 15%**. Power calculations use the sampling distribution from the original studies and the sample size estimations assume an equal amount of sampling from each treatment group.

**Figure S7: Power (A) and sample size simulations (B) for evenness for differentiating between non-obese versus obese for effect sizes of 1, 5, 10, and 15%**. Power calculations use the sampling distribution from the original studies and the sample size estimations assume an equal amount of sampling from each treatment group.

**Figure S8: Power (A) and sample size simulations (B) for the relative abundance of *Bacteroidetes* for differentiating between non-obese versus obese for effect sizes of 1, 5, 10, and 15%**. Power calculations use the sampling distribution from the original studies and the sample size estimations assume an equal amount of sampling from each treatment group.

**Figure S9: Power (A) and sample size simulations (B) for the relative abundance of *Firmicutes* for differentiating between non-obese versus obese for effect sizes of 1, 5, 10, and 15%**. Power calculations use the sampling distribution from the original studies and the sample size estimations assume an equal amount of sampling from each treatment group.

**Figure S10: Power (A) and sample size simulations (B) for relative risk of obesity based on Shannon diversity**. Power calculations use the sampling distribution from the original studies and the sample size estimations assume an equal amount of sampling from each treatment group.

